# Junior scientists spotlight social bonds in seminars for diversity, equity, and inclusion in STEM

**DOI:** 10.1101/2021.12.05.471284

**Authors:** Evan A. Boyle, Gabriela Goldberg, Jonathan C. Schmok, Jillybeth Burgado, Fabiana Izidro Layng, Hannah A. Grunwald, Kylie M. Balotin, Michael S. Cuoco, Keng-Chi Chang, Gertrude Ecklu-Mensah, Aleena K. S. Arakaki, Noorsher Ahmed, Ximena Garcia Arceo, Pratibha Jagannatha, Jonathan Pekar, Mallika Iyer, DASL Alliance, Gene W. Yeo

## Abstract

Disparities for women and minorities in science, technology, engineering, and math (STEM) careers have continued even amidst mounting evidence for the superior performance of diverse workforces. In response, we launched the Diversity and Science Lecture series, a cross-institutional platform where junior life scientists present their research and comment on equity, diversity, and inclusion in STEM. We characterize speaker representation from 79 profiles and investigate topic noteworthiness via quantitative content analysis of talk transcripts. Nearly every speaker discussed interpersonal support, and three-fifths of speakers commented on race or ethnicity. Other topics, such as sexual and gender minority identity, were less frequently addressed but highly salient to the speakers who mentioned them. We found that significantly co-occurring topics reflected not only conceptual similarity, such as terms for racial identities, but also intersectional significance, such as identifying as a Latina/Hispanic woman or Asian immigrant, and interactions between priorities and identities, including the heightened value of friendship to the LGBTQ community, which we reproduce using transcripts from an independent seminar series. Our approach to scholar profiles and talk transcripts serves as an example for transmuting hundreds of hours of scholarly discourse into rich datasets that can power computational audits of speaker diversity and illuminate speakers’ personal and professional priorities.

## Introduction

Following spring of 2020, broader recognition of widespread social injustice spurred advocacy for equity, diversity, and inclusion in STEM. Several teams of junior scientists have published recommendations on faculty hiring, grant review, and diversity initiatives ^1–4^. Irrespective of the disparity targeted, guidelines for equity, diversity, and inclusion build on a common foundation: listening to members of historically excluded groups share their experience working in STEM.

In June of 2020, we founded the Diversity and Science Lecture series (DASL), a platform where junior life scientists in San Diego and beyond can share their research and comment on equity, diversity, and inclusion in STEM (Figure 1a). DASL features integrated presentations on speakers’ personal backgrounds, their scientific progress, and their advice for navigating a scientific career. Each academic quarter, executive team members recruit speakers and schedule times for dry-run practice sessions and formal seminars. Each seminar consists of either two 15-20 minute talks from trainee life scientists or a 1 hour talk from an early career life scientist or social science expert.

**Figure 1:**
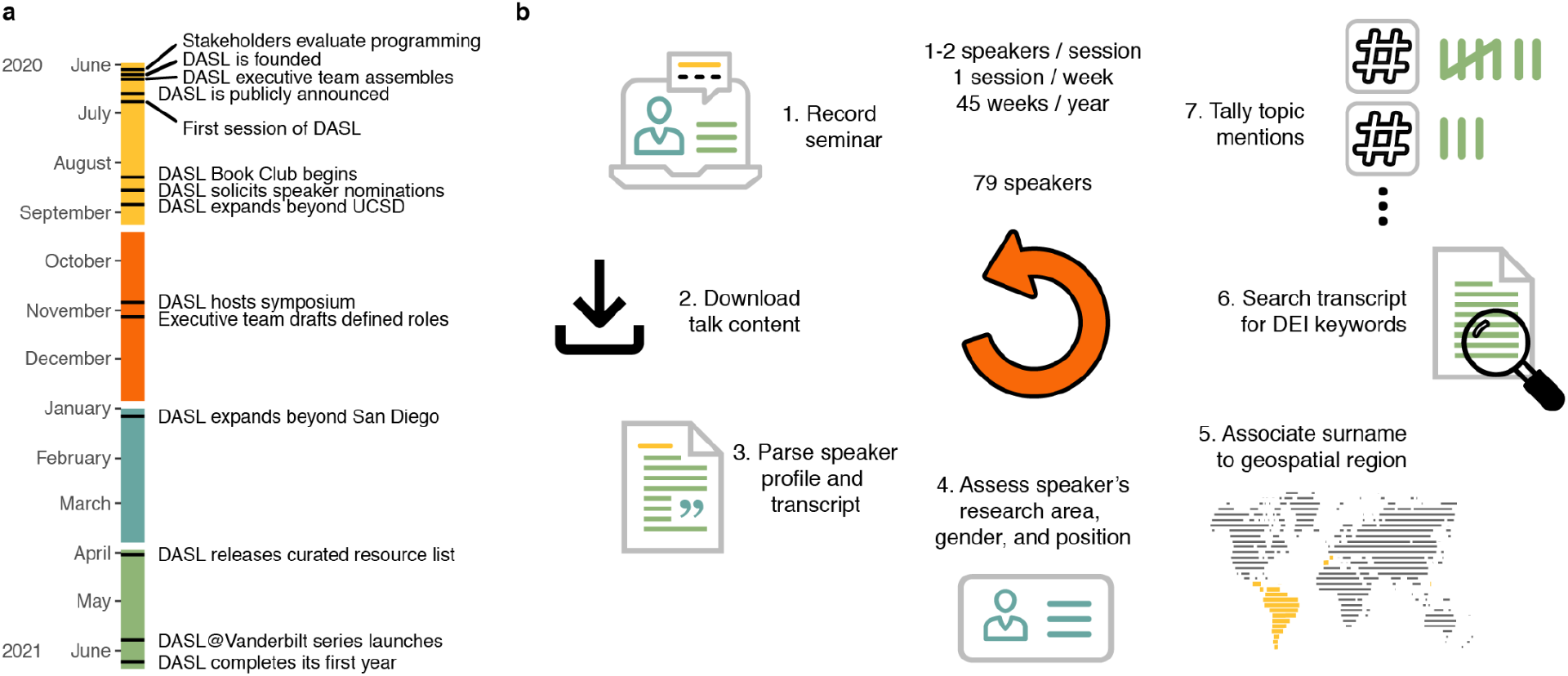
Reviewing the first year of DASL seminars. a) A timeline of DASL milestones separated by quarter. DASL was founded in June 2020 and completed its recent spring series in June 2021. b) Outline of the approach taken to synthesize insight from 79 weekly DASL seminar series speakers.

Other prominent virtual seminar series launched in 2020 commonly hewed close to existing scientific communities ^5–7^. By contrast, DASL is accessible to a broad cross-section of life scientists. Speakers are encouraged to speak freely about issues important to them and deliver a clear message advising their peers on how to advance equity, diversity, and inclusion in STEM. In June of 2021, DASL concluded a year of weekly sessions and continues to hold monthly sessions in its second year.

Who claims the opportunity afforded by DASL, and what are speakers’ priorities for equity, diversity, and inclusion in STEM?

Answering such questions from unstructured text drawn from dozens of seminars entails tremendous discretion. Coding speaker profiles and speech into quantitative data diminishes speakers’ qualitative experiences, but basing decisions on qualitative impressions makes effective prioritization of goals difficult and risks inaction on comparatively important but less salient concerns.

Reproducible statistical analysis for diversity requires special care. Only a few identities, such as nationality, geography, and gender, are commonly public information. Self-identification promises low rates of misclassification but still poses logistical challenges: what, if any, prefilled options to offer, and how to handle missing data, spelling errors, misunderstandings, and possible identifiability of participants if results are published. Programmatic and semi-automated methods that make inferences based on names or documents have the potential to curtail response bias, protect participant privacy, and rescue data lacking self-identification. Indeed, participant name is sufficient for scalable and reproducible (if uneven) inference of race, nationality, gender, and geographic associations ^8–11^. However, some identities, such as class, rural or agricultural background, first generation status, and sexual and gender minorities are almost wholly inaccessible without self-identification.

Here, we summarize how trainee life scientists conceive of equity, diversity, and inclusion in STEM (Figure 1b). We find large differences in representation compared to a traditional seminar series and identify differences in topic noteworthiness that depend on speaker background and priorities. We thus demonstrate how treating seminar series as data can yield surprising insight into who a seminar series includes and what they value.

## Results

In summer 2021, DASL concluded its first year of programming (Figure 1a). What once started as a community at UC San Diego grew to include scientists affiliated with universities and research institutes nationwide. Sustaining open and impactful conversation amongst graduate students, postdoctoral scholars, and faculty who had never met in person entailed immense planning and effort (Supplementary Figure 1a), but over its first year, DASL continuously grew its subscriber base (Supplementary Figure 1b), accrued over 4,000 unique website visitors, and achieved roughly 3,500 hours of engagement from attendees (Supplementary Figure 1c).

Over the course of a year, DASL hosted 79 speakers. First, we summarized the research focus of DASL speakers by parsing keywords in talk titles from speaker profiles. “Cell” was the most used term, at nine mentions, followed by “protein” and “regulate” at seven mentions, and “cancer”, “develop”, and “metabolism” at six. Other top words typically related to human cellular and molecular biology (Figure 2a). Next, we examined information about the speakers themselves. The majority of speakers were graduate students (53%), followed by postdoctoral scholars (34%), and assistant professors (6%) (Figure 2b). Overall, a greater percentage of speakers were women (67%) relative to UCSD life scientists broadly (58% of graduate students, 48% of postdocs, and 28% of tenure-track faculty; Figure 2c). By comparison, the UCSD Cellular & Molecular Medicine departmental virtual seminar series featured 42% women professors in 2021 (n=31). Overall, women were more likely to participate in DASL than men (p = 5e-3, binomial test).

**Figure 2:**
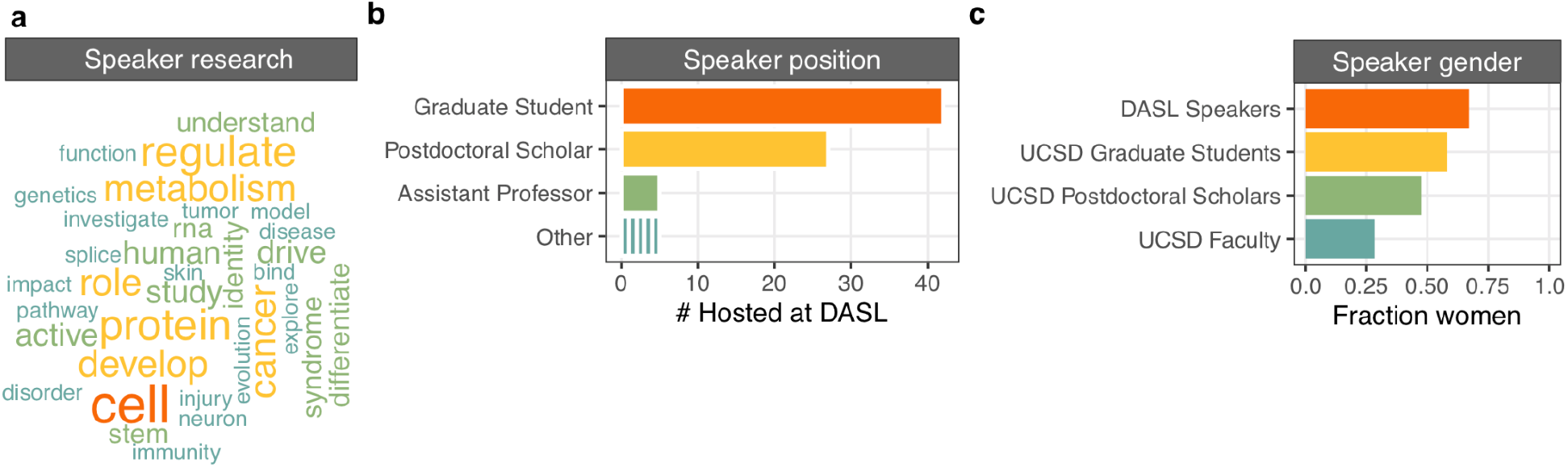
Analysis of DASL speaker profiles. a) Word cloud of the most common terms used in DASL research talk titles. The size of the word reflects the frequency. b) Breakdown of the positions held by DASL’s first 79 speakers. The ‘Other’ category includes Research scientist, Research Assistant Professor, Professor, Assistant Curator, and Administrator. c) Representation of women among DASL speakers compared to UCSD life science graduate students, postdoctoral scholars and faculty.

### Speaker surnames map a geography of DASL speaker diversity

To further characterize the diversity of speakers, we determined the country or region in the world where each speaker’s surname was most prevalent and abundant as reported by the Forebears database (see Methods). For most names, prevalence and abundance pointed to the same region, enabling imperfect but scalable inference of where relatives of speakers currently reside. (Figure 3a). To reflect historical global migrations and corresponding name usage, names linked to Britain, the United States, Canada, Australia, New Zealand and the Caribbean were grouped in a separate Anglophone category – surnames that were most abundant in Anglophone countries but more prevalent in a non-Anglophone region were annotated in accordance with the non-Anglophone region.

**Figure 3:**
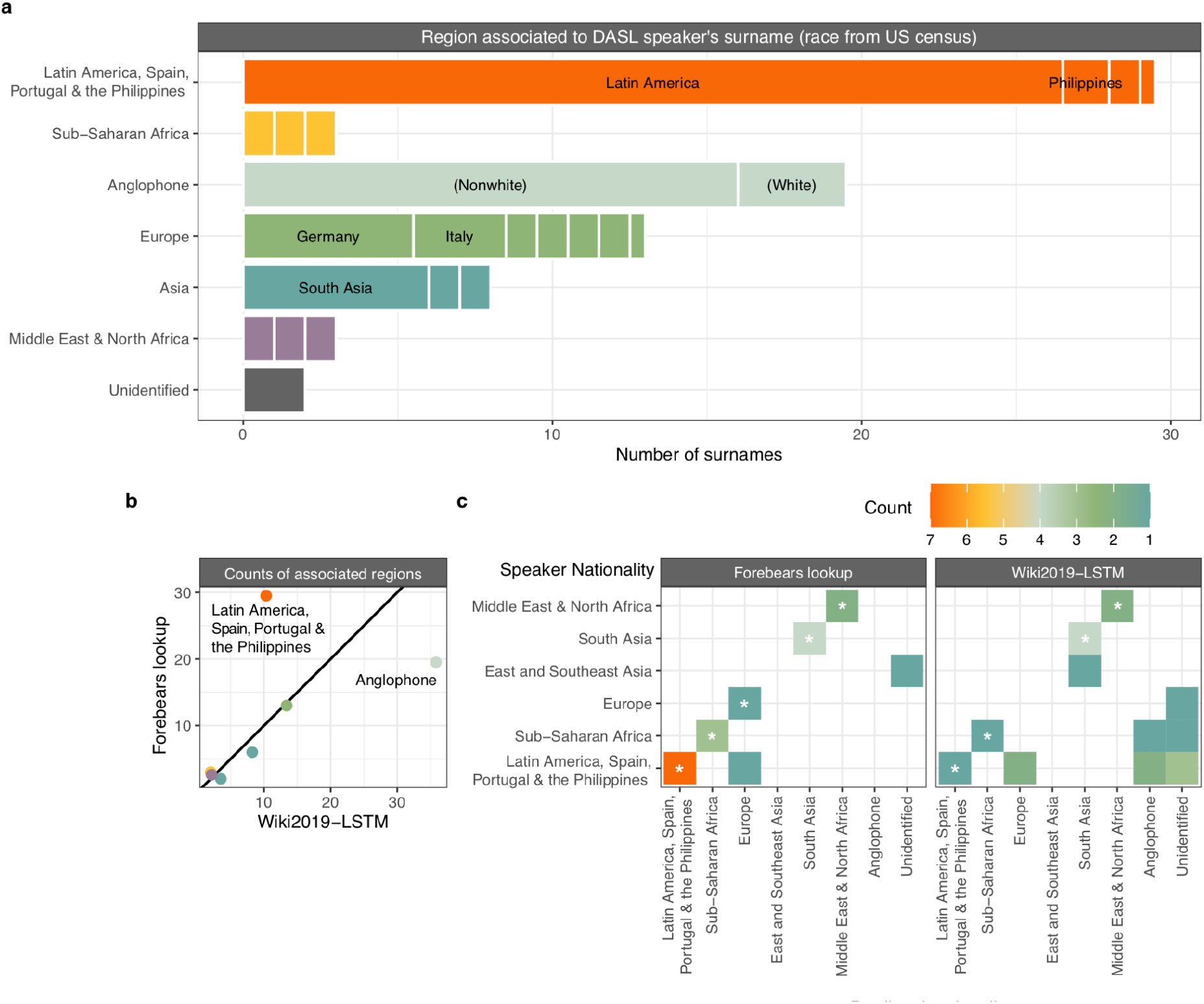
Association of DASL speaker names to geospatial groups. a) Counts of most associated regions for DASL speakers’ surnames using the geospatial name database Forebears. ‘Anglophone’ refers to names associated to Britain, the United States, Canada, Australia, New Zealand or a country in the Caribbean. b) Aggregate counts for associated regions using Wiki2019-LSTM (x-axis) and the Forebears database (y-axis) across all DASL speakers. c) Confusion matrices for international DASL speaker names (19 surnames from 14 speakers) demonstrating accuracy of predictions for speaker nationality. Asterisks denote correct identifications. Names for which Wiki2019-STLM achieved less than 50% probability for a region were labeled as “Unidentified”.

Overall, Latin American (30 names) countries were the top associated regions for DASL speakers’ surnames. Next most common were Anglophone (20 names) and European countries (13 names) (Figure 3a). We further annotated anglophone-associated surnames as “White” or “Nonwhite” using US Census data linking common US surnames to race ^10^. Using this approach, 82% of Anglophone speaker surnames were labeled “Nonwhite” in DASL.

We performed the same analysis for 63 speakers hosted by the Fragile Nucleosome forum, a concurrent international remote lecture series that has promoted community building (Supplementary Figure 2) ^7^. 30 Fragile Nucleosome speaker names were associated to European countries and 14 to Anglophone countries. Most Anglophone surnames (64%, n=14) were labeled “White.” Overall, 30% of Fragile Nucleosome speaker surnames were associated to regions outside Europe and Anglophone regions compared to 58% for DASL speaker surnames.

We next compared lookup of surname-associated regions in Forebears to imputation using a recently published long short-term memory recurrent neural network trained on names and nationalities scraped from Wikipedia (“Wiki2019-LSTM”) ^8^. For most geographic regions, predictions agreed well in aggregate, but Anglophone names and names associated to Latin America, Spain, Portugal, and the Philippines were discrepant (Figure 3b). We re-examined predictions for 14 speakers who publicly identified their nationality and found a 71% success rate for lookup in Forebears versus 36% for LSTM predictions. LSTM predictions for Latin American speakers’ names were often linked to Anglophone regions (Figure 3c), suggesting that overrepresentation of contributors from Anglophone countries in Wikipedia may cause bias against Latin American names in the learned LSTM.

### DASL speakers emphasize the importance of social and interpersonal factors in STEM

DASL speakers covered a broad range of topics: childhood familiarity with a career in STEM, the burden of fees in graduate admissions, immigrant identity, coming and being out as queer or trans in academia, navigating parenthood in STEM, mental health challenges in academia, health disparities for racial minorities, advocacy for people with disabilities, the complexity of multi-racial identity, cultural expectations clashing with career pressure, ethical research for indigenous communities, anti-science attitudes in rural America, and both subtle and overt racism against Black people and people of Middle East and North Africa descent.

By counting mentions of keywords for each trainee DASL talk, we tracked common themes (Figure 4a). Topics related to social factors were most likely to be broadly discussed. “Education” keywords were most frequently mentioned (46/54 talks), followed by “Family” keywords (39/54) and “Mentoring” keywords (37/54). Keywords for the concepts “Collaboration” and “Success” were also mentioned by a majority of speakers, but typically received few mentions per speaker. Other topics were mentioned many times by a small number of speakers: “Sexual and gender minorities”, “Mental health”, and “Finances”. Mentions of “Race/ethnicity” and “Friends” were intermediate in both regards: speakers broke roughly evenly between zero mentions, one or two mentions, or many mentions. Thus, topics’ noteworthiness varied considerably both in breadth (fraction of speakers mentioning a topic) and salience (the typical number of mentions per speaker).

**Figure 4:**
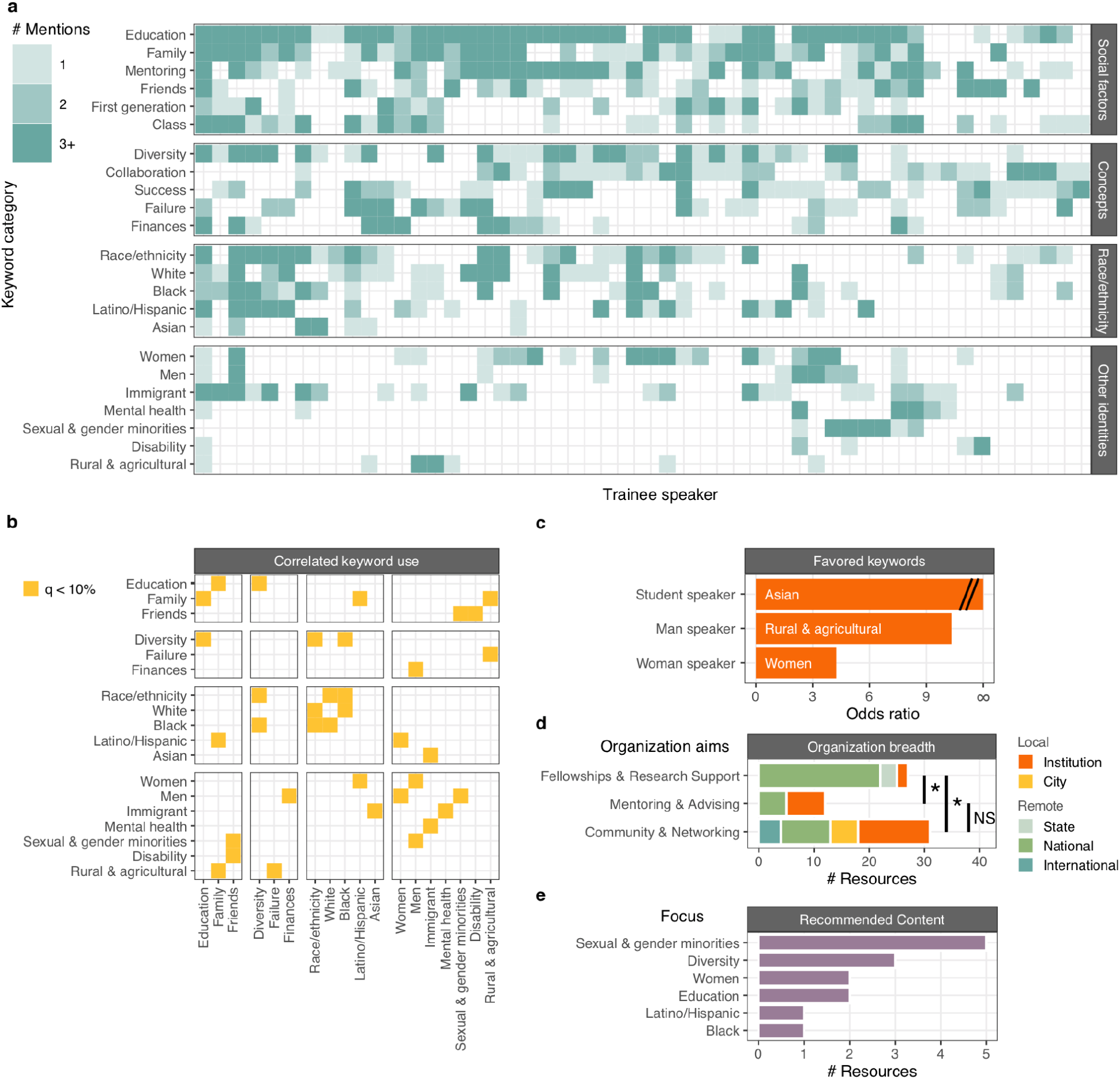
Topics relevant to equity, diversity, and inclusion mentioned by trainee DASL speakers. a) Truncated counts (more than 3 mentions were placed in “3+”) of mentions of topics across speakers (x-axis) organized by topic (y-axis) and broader type of concern (social, race/ethnicity, or other identities). b) Significantly correlated keyword usage for terms in a). c) Differential keyword use stratified by speaker identity. d) Helpful organizations cited by DASL speakers, annotated by organization aims and scope. e) Count of DASL speakers’ recommended content stratified by focus.

### Topics discussed vary by speaker identity and intersect with one another

18 pairs of topics exhibited correlated keyword usage (by permutation of spearman correlation coefficient, 10% FDR) (Figure 4b). The most significantly correlated topics were “Diversity” and “Education” (rho = 0.465, p = 4e-4). Whereas “Friends” correlated with “Sexual & gender minorities” and “Disability”, “Family” correlated with “Latino/Hispanic” and “Rural & agricultural”. Indeed, 100% of talks mentioning “Latino/Hispanic” keywords also mentioned family, strongly linking discussion of Latino and Hispanic identity to family values. Conversely, disability and sexual and gender minority identities were linked to making friendship a priority.

Other correlated themes supported nuanced interpretation of the data. Mentions of “Immigrant” correlated with “Asian” but not other racial groups, and “Latino/Hispanic” with “Women” but not men, suggesting the existence of special intersectional identities. Similarly, mentions of “Sexual & gender minorities” and “Finances” were correlated with “Men” but not “Women” keywords.

In rare cases, the likelihood of a topic depended on the degree status or gender of the speaker (Figure 4c). Men speakers were 10 times more likely to mention “Rural & agricultural” keywords than women speakers (Binomial test, p = 3e-4), women speakers were 4 times more likely to mention “Women” keywords than men speakers (Binomial test, p = 4e-4), and graduate student speakers were more likely to mention “Asian” keywords than postdoctoral scholars (Binomial test, p = 2e-3) (10% FDR). We expect that smaller but still significant differences in topic mentions across speaker identities could be detected with a larger pool of speakers.

### DEI speakers’ focus on social and interpersonal factors persists across seminar series

To learn more about the trends we identified (Figure 5a), we applied the same keyword analysis approach to another UC San Diego DEI seminar series: Scientific Queers United in Academic Discourse (SQUAD). In contrast to DASL, SQUAD features approximately 40-minute talks exclusively from LBGTQ+ scientists based in San Diego and does not seek speakers from a specific career stage or discipline. SQUAD speakers do address both their research and their personal experiences, as in DASL seminars.

**Figure 5:**
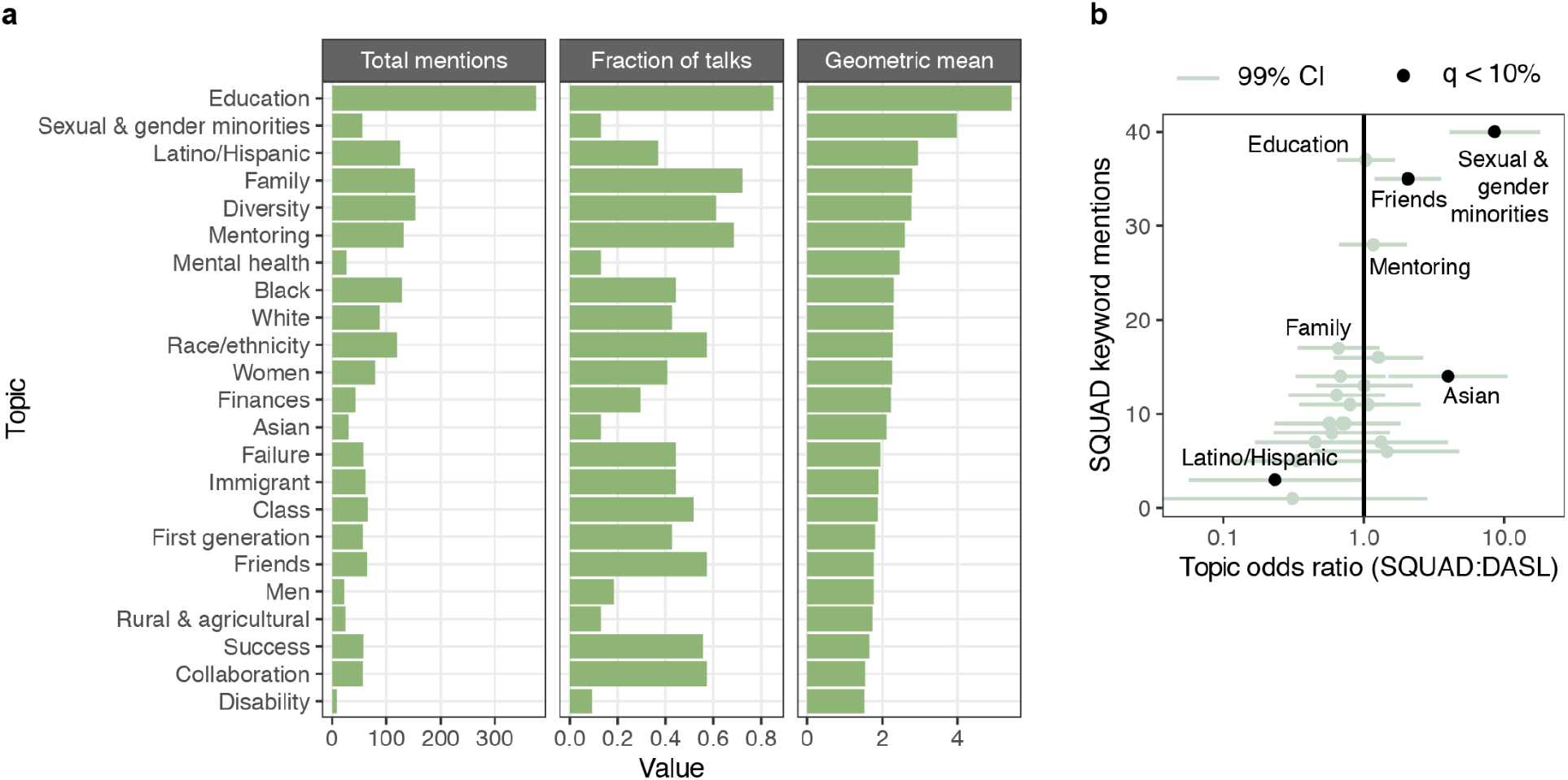
Quantifying and contextualizing topic noteworthiness at DASL. a) Count of the total mentions per topic (left), topic breadth, or the fraction of talks with at least one mention (middle), and topic salience, or the geometric mean of the number of mentions per talk with at least one mention (right). b) Comparison of DEI topic mentions in DASL to SQUAD by odds ratio. Each topic mentioned by a SQUAD speaker and its 99% confidence interval are plotted.

Unsurprisingly, SQUAD speakers most often mentioned “Sexual & gender minorities” keywords - mentions were 8.5 times more likely than in DASL seminars (Figure 5b). Otherwise, SQUAD speakers largely prioritized the same social and interpersonal factors highlighted by DASL speakers. The next most frequent topics at SQUAD after “Sexual & gender minorities” were, in order, “Education”, “Friends”, “Mentoring”, and “Family” (Figure 5b).

We next assessed more subtle differences in keyword usage between DASL and SQUAD. SQUAD speakers were 4 times more likely to mention “Asian” keywords (Binomial test p = 5e-4), 2.1 times more likely to mention “Friends” keywords (Binomial test p = 1e-3), and 4.3 times less likely to mention “Latino/Hispanic” keywords (Binomial test p = 5e-3), after applying a 10% FDR (Figure 5b). The greater use of “Friends” keywords in SQUAD seminars echoes our earlier observation of correlated usage of “Sexual & gender minorities” and “Friends” keywords. Notably, SQUAD speakers did not uniformly increase mentions of social factors: family keyword use was less frequent, if anything (p > 0.05). Transcripts of SQUAD seminars are thus further evidence of greater relative attention paid to friendship among sexual and gender minorities.

### Speaker’s resources suggest social bonds are vulnerable to disruption by social distancing

In addition to keyword analysis, we curated specific resources DASL speakers cited as helpful to their scientific careers. The 90 entries include entities offering fellowship and research support, community and networking groups, mentoring and advising centers, professional societies, and recommended content ^12^. Cited mentoring and advising entries (Fisher exact test, p = 6e-5) and community and networking entries (Fisher exact test, p = 1e-3) were 16 times more likely to operate at local levels compared to fellowship and research support opportunities (Fisher exact test, p > 0.05) (Figure 4d). While funding sources may be fungible and robust to social distancing, we infer that community-building and mentoring opportunities that trainee scientists relied on were greatly diminished by pandemic precautions.

For the 14 recommended websites and articles (“Recommended content”), we labeled each with the most relevant topics describing their content. The plurality (5/14) commented on sexual and gender minorities, followed by diversity broadly (3/14) and women and education (each with 2/14).

## Discussion

In this work, we examined 79 speaker profiles and 54 talk transcripts from the Diversity and Science Lecture series. We characterized who spoke at DASL and the topics they discussed using a quantitative content analysis framework. We cross-referenced the gender balance of DASL speakers against that of UCSD life science trainees and found greater participation from women scientists. We also exploited a geospatial analysis of speaker surnames that showed particular enthusiasm from Latino and Hispanic scientists. Our tallies of keyword mentions in talk transcripts indicated that social factors like family, education, and mentorship are highly salient and broadly noteworthy (Figure 5a). Sexual and gender minority was noteworthy to fewer life science trainees but still highly salient. Our review of content recommended by speakers addressed offered more evidence for sexual and gender minority identity to select trainees (Figure 4e).

It is perhaps no surprise that unprecedented social distancing measures prompted trainee life scientists to reflect on the contribution of interpersonal relationships to equity, diversity, and inclusion in STEM. Yet, that discussion of social factors including family and mentorship surpassed that of racial identity in a diversity-themed seminar series deserves note. Academic literature on diversity broadly, beyond the purview of trainee life scientists, addresses race and ethnicity (9750 articles in a Google Scholar search) much more often than family (4040 articles) or mentoring (445 articles). Furthermore, the noteworthiness of social factors like education, family, and mentoring stood out not only because nearly all speakers addressed them, but because each speaker commented on them repeatedly (Figure 5). We believe that trainee life scientists highly esteem a sense of camaraderie and that access to interpersonal support is essential for achieving equity, diversity, and inclusion at academic institutions.

Our correlational analysis suggests special care be taken regarding intersectional identities and common needs within historically excluded groups. Particularly in light of recent anti-Asian sentiment in North America, Asian immigrant identity comes with unique burdens that cannot be recapitulated from separate Asian and immigrant identities. Similarly, Latina women face specific stereotypes that do not burden women or Latino/Hispanic people broadly. The social ties highlighted by speakers also depended on the identities discussed. Friends and family were each mentioned by a majority of speakers, but friendship garnered more commentary in talks mentioning sexual and gender minorities and disability, and family garnered more commentary in talks mentioning Latino/Hispanic and rural background. It is reasonable to infer that sexual and gender minorities and people with disabilities on average especially prioritize friendship in order to connect with those with a shared identity. Similarly, Latino/Hispanic individuals and those from rural backgrounds especially prioritize family. Thus, disruption of time spent with either friends or family would be expected to have disparate impacts on these groups. We advise greater awareness of how social distancing measures might affect historically excluded groups differently and to invest in approaches to mitigation accordingly.

The methods we present do not solve all challenges regarding data on diversity in STEM. Aggregating identities with divergent lived experiences into broad categories (“Asian” or “Sexual and gender minorities”) can mask important disparities for subgroups (Southeast Asian or transgender identity), but low counts for more rarely adopted or hard-to-define identities preclude statistical inference. Indeed, multi-racial identity was not examined for this reason despite vigorous discussion of this topic in DASL sessions. Language describing identity can rise and fall in popular usage and complicate comparison to data collected even a few years in the past or future. Finally, balancing topics of high salience to a small subset (e.g., sexual and gender minorities) against topics of modest salience to a majority (e.g., the role of collaboration in research) is not easy, but necessary to realize an inclusive environment in STEM.

Past studies have suggested that virtual events can achieve greater equity and inclusion for junior career scientists in diverse cultural and institutional settings ^13,14^. We believe that the greater visibility for junior life scientists made possible by DASL is beneficial in its own right. Speakers reported unanimous approval for dry runs that allowed them to receive feedback on their presentations before formal sessions. Both formal seminars and meet-the-speaker sessions allowed speakers to network with other scientists, although we report that meetings scheduled after seminars achieve lesser engagement than dry runs. With this in mind, it is worth questioning why some identities were poorly represented. Men and Asian DASL speakers - especially Asian men speakers - were few. Discussion of Asian identity occurred explicitly only in graduate student talks. Topics specifically relevant to men speakers included finances, sexual and gender minorities, and rural and agricultural background. The full implications of these trends are unclear, but we urge awareness of how Asian and men scientists might have different priorities for diversity initiatives that go beyond discussion of race and gender.

Attention is a fiercely contested resource in professional settings; that trainee life scientists directed the attention of hundreds of their peers and mentors stands as an undeniable achievement. Nonetheless, disparities for women ^15–19^, racial minorities ^8,20–23^, and sexual and gender minorities ^24–27^ working in scientific disciplines have been well documented and urgently demand further study and remediation. Recruiting dozens of life scientists for a public seminar series is both more practical and enriching for the scientific community than conducting a study based on interviews or focus groups and more sustainable than regularly administering surveys. Furthermore, as we demonstrated, publicly accessible seminar series can be parsed to provide additional context or offer insight into the priorities of STEM workers without hosting a seminar series at all. We believe that harnessing speaker profiles and transcripts from scientific seminars for content analysis will prove broadly useful for conducting metascience research, and especially for characterizing the priorities of women and minorities in STEM.

## Methods

Representation of women was computed from UCSD diversity dashboards. Graduate students in the Health Sciences and Biological Sciences who chose “Woman” as their gender were counted. For postdocs, academic personnel with the appointment title “Postdoctoral scholar/fellow” in the Health Sciences and Biological Sciences were counted. For faculty, the “ladder-rank professor” appointment title was selected. Data was taken from fall 2019, the most recent data available. Probability of success in null binomial testing was weighted by the fraction of women at each career stage (50.7% women overall).

Terms in talk titles were counted by parsing titles posted on the DASL website. Text was tokenized by tidytext::unnest_tokens and stop words removed using the tidytext’s stop_words dataset. Words were stemmed using SnowballC::wordstem, and the number of titles containing each stemmed word were counted. The frequency of retained stems that occurred in at least 3 titles were converted back into representative words and visualized using the wordcloud package.

Speaker surnames were noted from the DASL website and Fragile Nucleosome website over the same time period. Surnames were associated to geographic regions using Forebears.io, the “largest geospatial names database”, which purportedly aggregates records from over 27 million surnames and 4 billion individual records from 236 countries or jurisdictions^28^. Jurisdictions in Latin America were pooled into one region. Jurisdictions in Britain, the United States, Canada, Australia, New Zealand and the Caribbean were pooled into an Anglophone region because of their shared British surnames ^29,30^. South Asian jurisdictions were pooled into a South Asian region. For each surname, the region with the highest incidence and frequency were noted. Names from speakers with two surnames were weighted accordingly. For surnames that matched multiple regions, the most frequent region only applied when the incidence was above 500.

Surnames were considered validated 1) if the most prevalent region was also the most frequent, 2) if the most prevalent region was at least tenfold greater than the most frequent region (most prevalent region chosen), 3) if the most prevalent region was the Anglophone region and less than tenfold greater than the most frequent region (most frequent region chosen). If these conditions were not met, or there was no match to the database, the surname was deemed unidentified. Surnames assigned to the Anglophone region were further annotated with race using the predictrace::predict_race command in R. Surnames assigned high confidence for matching White US Census takers were labeled “(White)”. Surnames that were not high confidence for corresponding to White US Census takers were labeled “(Nonwhite)”. Note that these methods are not sensitive for Native American and Alaskan Native or multiracial identities reported to the US Census^31^.

The Wiki2019-LSTM model was downloaded from its GitHub repository (https://github.com/greenelab/wiki-nationality-estimate). The 07.test-ismb-data.py script was modified to issue predictions on DASL speaker surnames. For aggregate counts, fractional assignments were summed across all speakers. For visualization of the confusion matrix, surnames that had no region above 50% probability were labeled as “Unidentified”. For the sake of comparison with Forebears lookup, the Wiki2019-LSTM’s Celtic/English category was deemed synonymous with our Anglophone category, and the Wiki2019-LSTM’s Israel category was pooled with our Middle East and North Africa category.

Publicly viewable DASL and SQUAD talks were downloaded and uploaded to YouTube to obtain transcripts. Talks were reviewed and keywords tallied in R. Exact keyword expressions and counts are available in the supplementary data. Mentions of “white matter” and other brain science terms were manually removed from counts of “White” mentions. Column order was determined by running the R command hclust on Pearson correlation distance for a matrix of the log10 of counts plus 1 which was then scaled with the scale command for each keyword across 54 talks. The number of mentions of each keyword were truncated at 3 or more for data visualization purposes. Significantly correlating terms were determined using the R command cor(m=”s”) on the matrix of total mentions by speaker and keyword and removing duplicate comparisons; p-values were computed from an empirical distribution using the ecdf command on permuted data. Differential mentions by speaker status and seminar series were determined by using the R command binom.test on the sum of truncated mentions (DASL speakers with more than 3 mentions were replaced by 3 and SQUAD speakers with more than 6 replaced by 6 to mitigate the impact of outliers) split by gender, degree status or seminar series. The false discovery rate was set using the p.adjust command. Binomial confidence intervals were obtained using the binconf command from the Hmisc package.

The Google Scholar search queries “allintitle: diversity (race OR racial OR ethnic OR color)”, “allintitle: diversity (mother OR mothers OR father OR fathers OR son OR sons OR family OR families)”. and “allintitle: diversity (mentor OR mentors OR mentorship OR mentoring)” were used to estimate the relative focus of diversity literature on race, family and mentoring.

Icons for the DASL schematics were downloaded from The Noun Project (nounproject.com). Lightbulb by Maxim Kulikov, Rules by Adrien Coquet, Webinar by ProSymbols, Whisper by ProSymbols, Toolbox by WEBTECHOPS LLP, Document by Jamison Wieser, downloads by Gregor Cresnar, Count by Deylotus Creative Design, redo by Creative Stall, Magnifying Glass by Chatchai Pripimuk, World Map by Roussy lucas, hastag by Tomi Triyana, identity by Saurus Icon.

## Acknowledgements

DASL is funded in part by a grant from the Chan-Zuckerberg Initiative. E.A.B. acknowledges funding from the Helen Hay Whitney Foundation. G.W.Y. is an Allen Distinguished Investigator, a Paul G. Allen Frontiers Group advised program of the Paul G. Allen Family Foundation. The 2020 DASL symposium was supported by HHMI Gilliam Fellows, the ThermoFisher Queer Working Group, the Huntsman Cancer Institute, the UCSD Biology Student Social Fund, UCSD Graduate Division, UCSD Biomedical Sciences Graduate Program, the UCSD Biology Diversity Committee, the UCSD Office for Equity, Diversity, and Inclusion, the UCSD Latinx/Chicanx Academic Excellence Initiative, the UCSD Black Academic Excellence Initiative, and the UCSD School of Medicine. We thank the oSTEM UCSD graduate chapter leadership for hosting SQUAD and making seminars available on their website.

## Supplementary materials

Supplementary Data: DASL Alliance author list, nationality validation data, and search queries and keyword counts for talk transcripts

---

**Supplementary Figure 1:**
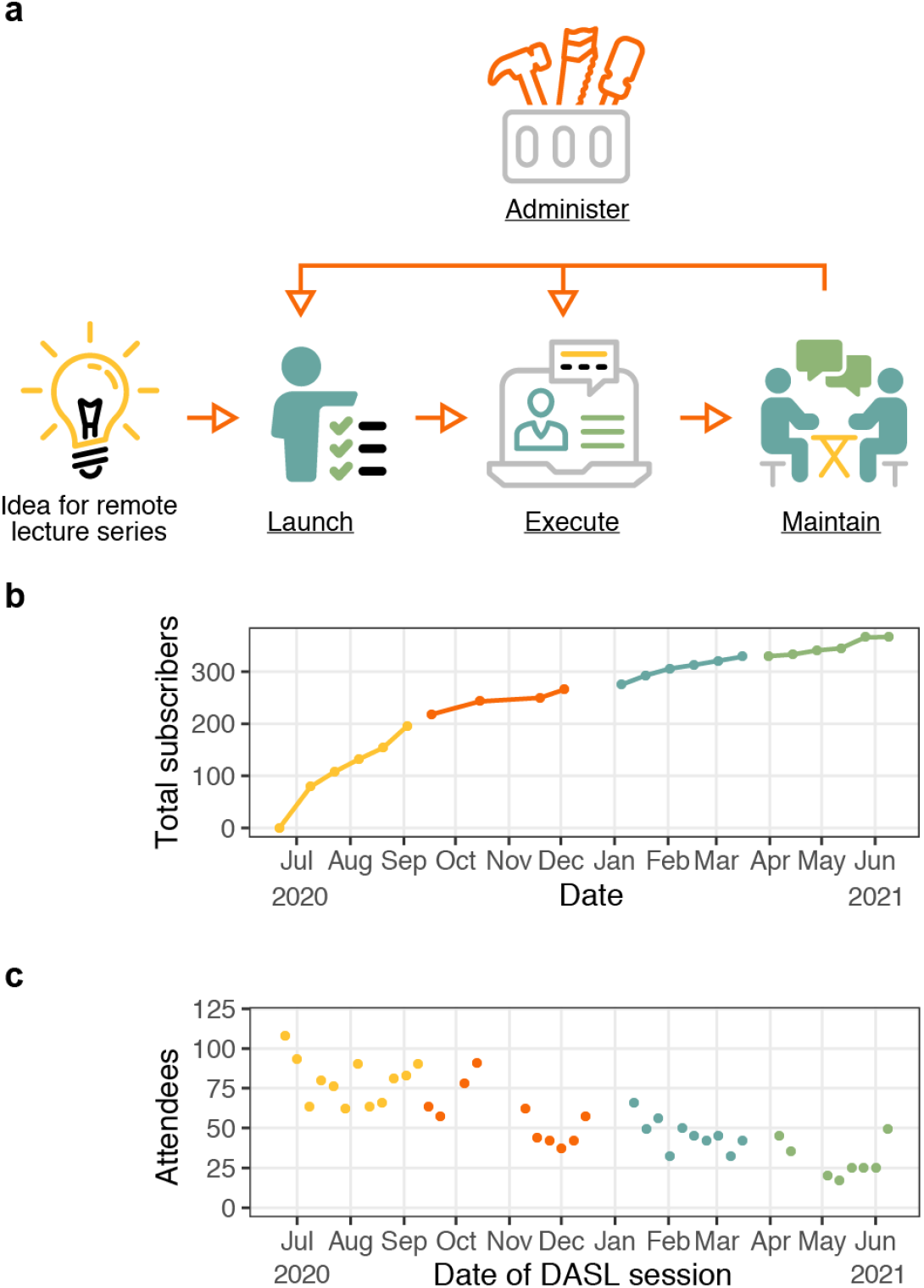
Engagement in the first year of DASL. a) Outline of running a remote lecture series. We provide guides detailing how to approach each underlined component on our website. b) Total subscribers to the DASL mailing list over time. c) The number of attendees per DASL session over the first year of programming.

**Supplementary Figure 2:**
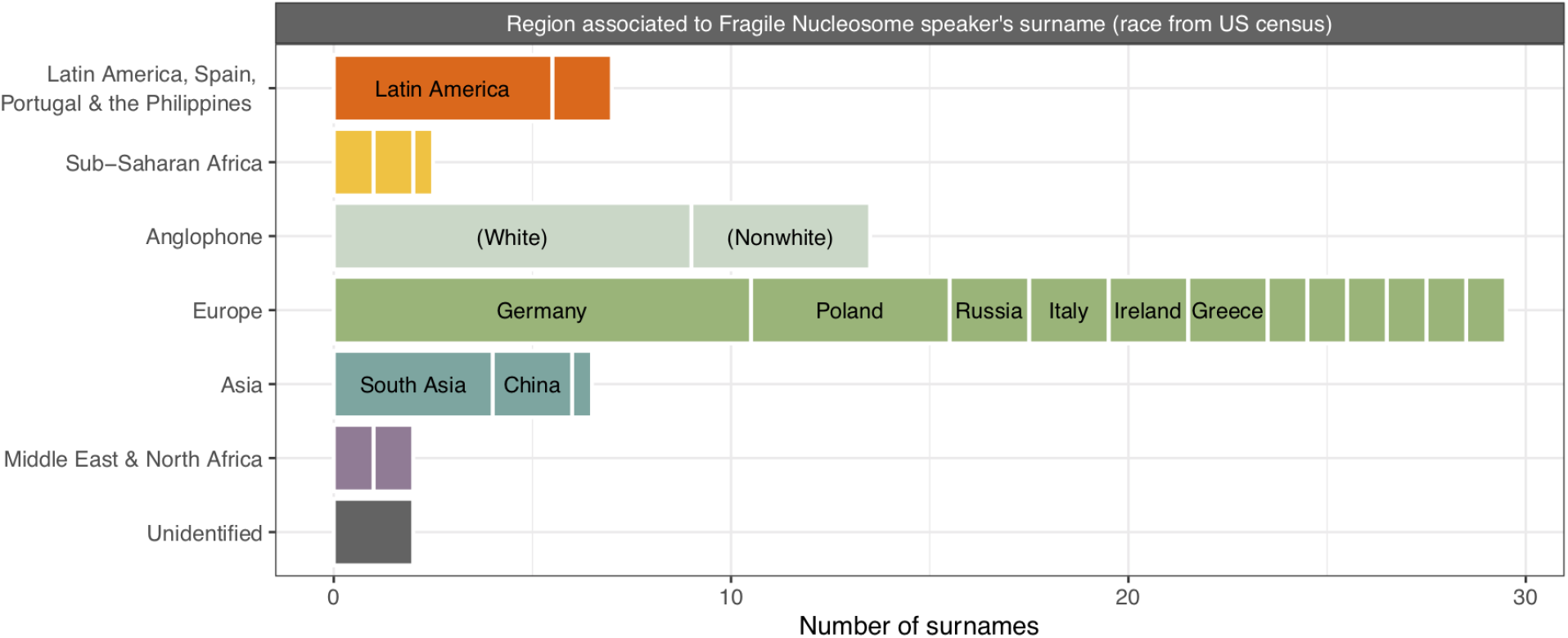
Support for surname associations. Speaker name associations from the Fragile Nucleosome seminar series. Counts are tallied as in Figure 3a.

